# Copper reduces the virulence of bacterial communities at environmentally relevant concentrations

**DOI:** 10.1101/2023.06.02.543412

**Authors:** Luke Lear, Dan Padfield, Elze Hesse, Suzanne Kay, Angus Buckling, Michiel Vos

**Author notes:** Corresponding author;. Environment and Sustainability Institute, University of Exeter, Penryn, Cornwall, United Kingdom, TR10 9FE.

## Abstract

Increasing environmental concentrations of metals as a result of anthropogenic pollution are significantly changing many microbial communities. While there is evidence metal pollution can result in increased antibiotic resistance, the effects of metal pollution on virulence remains largely undetermined. Here, we experimentally test whether metal stress alters the virulence of bacterial communities. We do this by incubating three wastewater influent communities under different environmentally relevant copper concentrations for three days. We then quantify the virulence of the community using the *Galleria mellonella* infection model, and test if differences are due to changes in the rate of biomass accumulation (productivity), copper resistance, or community composition (quantified using 16S amplicon sequencing). The virulence of the communities was found to be reduced by the highest copper concentration, but not to be affected by the lower concentration. As well as reduced virulence, communities exposed to the highest copper concentration were less diverse and had lower productivity. This work highlights that metal pollution may decrease virulence in bacterial communities, but at a cost to diversity and productivity.

## Introduction

Many metal ions are essential for metabolism and consequently for life (1). However, what gives metals their beneficial properties – their ability to cycle between oxidation states – also causes them to become toxic at high concentrations (1, 2). It is therefore essential for organisms to finely balance cellular concentrations of metal ions in order to survive (3). This balance may become increasingly harder to achieve in anthropogenically polluted environments, where microbes previously unexposed to metal stress are rapidly exposed to potentially toxic concentrations of metal ions (4–10).

Metal pollution can reduce the microbial diversity, productivity, biomass, and function of bacterial communities (8, 11, 12). These effects in turn can be detrimental to human health by reducing soil fertility and decomposition of organic matter (8). Furthermore, many common metal detoxification mechanisms used by bacteria, such as efflux pumps and reduced cell membrane permeability (13, 14), can pose an indirect threat to human health by conferring resistance to antibiotics (13, 15). However, whilst there is strong evidence for increased antibiotic resistance in metal-polluted environments (16), the effect of metals on bacterial virulence remains largely unknown. This is an important knowledge gap because simultaneous selection for antibiotic resistance and virulence could significantly exacerbate disease with detrimental effects on human health.

There are several possible ways in which bacteria could increase both metal resistance and virulence, including biofilm formation (17–19), efflux pumps (20) and the production of metal-sequestering siderophores (21). However, to date only siderophores have been directly linked to both virulence and metal detoxification (21). Siderophore production, in addition to being a metal detoxification mechanism, aids iron uptake – an essential nutrient for microbial growth that is commonly of limited availability within hosts (20). Consequently, upregulation of siderophore production in response to copper can increase virulence by increasing within-host growth, as recently shown in experimental populations of *Pseudomonas aeruginosa* (21). Like siderophores, increased biofilm production is positively associated with virulence, as it can enable greater population densities by facilitating resistance to host defences and adherence to host cells (22). However, despite evidence that metals can select for biofilm production because they can form a protective layer due to their anionic properties binding and trapping cationic metal ions (17), it remains unknown whether metals can select for virulence through this mechanism. In contrast to siderophore and biofilm production, efflux pumps are both positively and negatively associated with virulence (see Fernando *et al,* 2013 for a comprehensive review (20)). Positive associations have been found when pumps increase resistance to host-derived antimicrobials and aid toxin production (20), whereas negative associations have been found when overexpression of pumps leads to downregulation of certain virulence factors, such as Type III secretion proteins in *P. aeruginosa* (23).

Previous studies on the effects of metals on virulence have mainly focused on changes in virulence at the phenotypic and genotypic level (*e.g.* (21)), and have not considered community-wide responses. Whilst understanding the links between specific metal resistance mechanisms and virulence within individual species is important, it is possible that metals may affect virulence differently at the community level. For example, siderophore production evolves differently at the community level compared to in single species populations (24). It is therefore possible that metals could select for or against virulence at the community level by changing the frequency of virulence traits within the community, either by affecting expression, *de novo* evolution, or species carrying these traits (species sorting). Moreover, as metals can reduce the diversity of bacterial communities (25, 26), effects of species sorting might be more important than physiological responses or evolutionary changes within individual species. Metal-induced richness and diversity losses may lead to virulent species being removed either selectively or stochastically from the community, or conversely may select for the most virulent species when metal detoxification traits also function to increase virulence. Furthermore, a change in diversity may subsequently affect the productivity of a community (27), which could result in a decrease in virulence as decreased growth rate can result in decreased acquisition of host resources (28–30).

Here, we experimentally test for the effects of copper pollution on the virulence of microbial communities by incubating microbial communities taken from wastewater influent with two copper concentrations at levels similar to that observed in metal-polluted environments, including water (31) and agricultural soils (32). We chose wastewater influent communities as they are likely to contain bacteria relevant to human health, and often contain species that can survive in the human microbiome and the wider environment, such as *Escherichia coli* and *Klebsiella pneumoniae* (33). To determine whether the effect of copper on virulence is consistent across multiple communities, we use three different wastewater samples, each taken from the same wastewater treatment plant, but in different years. The communities were incubated for three days (with daily transfers) with no, low, or high concentrations of copper, before being transferred into a common garden environment (copper-free media) for 24 hours to remove any phenotypic effects of copper (21). We then quantified virulence using an insect infection model, community productivity using growth rate data, and diversity using 16S amplicon sequencing.

## Methods

### Source of microbial communities

Three samples of raw wastewater influent (communities ‘A’, ‘B’ and ‘C’) were collected from a wastewater treatment plant in Falmouth, UK, in 2019, 2020 and 2021. Each sample was frozen in glycerol at a final concentration of 20% and stored at -80°C.

### Experimental design

Each of the three wastewater communities were inoculated into 18 separate 35mL glass vial microcosms (54 in total) containing 6mL of Iso-Sensitest broth (Oxoid, Basingstoke, United Kingdom) and incubated statically at 37°C. Six microcosms per community contained no copper, six contained copper sulphate (CuSO_4_; Alfa Aesar, Massachusetts, United States) at a concentration of 0.1g/L and six contained CuSO_4_ at a concentration of 1.0g/L. After 24 hours, each microcosm was thoroughly homogenised by vortexing and 60µL (1% of the volume) was transferred into fresh media of the same copper concentration. Transfers were carried out every 24 hours for three days. To control for the physiological effects of copper, on day three all microcosms were homogenised and 60µL transferred into Iso-Sensitest broth containing no copper (*i.e.* a common garden environment) (21). These cultures were grown for a further 24 hours before multiple aliquots were frozen at -70°C, both with glycerol at a final concentration of 25% (for use in virulence, growth rate assays and resistance assays) and without glycerol (for amplicon sequencing).

### *Galleria mellonella* virulence assay

Virulence of the passaged communities was quantified using the insect infection model *Galleria mellonella* (34), a well-established proxy for the mammalian innate immune system (28, 35). Furthermore, this model has recently been used to quantify virulence at the community level by injecting whole community samples (36–38). We first quantified the final density of each culture by plating defrosted samples onto Iso-Sensitest agar and counting the number of colony forming units (CFU) after 24 hours incubation at 37°C. We then used these counts to standardise each defrosted sample to 10^5^ CFU in M9 salt buffer, before 10µL was injected into 20 final instar larvae per replicate using a 50µL Hamilton syringe (Hamilton, Nevada, USA). Larvae were then incubated at 37°C and mortality checked every hour from 12 to 24 hours post-injection. Larvae were classed as dead when mechanical stimulation of the head caused no response. M9-injected and non-injected controls were used to confirm mortality was not due to injection trauma or background *G. mellonella* mortality; >10% control death was the threshold for re-injecting (no occurrences).

### Growth curves and quantification of copper resistance

The growth characteristics and resistance to copper of all 54 final communities was quantified using optical density measurements. Firstly, 20µL of each final community was transferred into three wells containing 180µL of Iso-Sensitest broth in a 96-well plate (total of three 96-well plates: one for each starting community). One well contained media with no copper, one contained copper sulphate at a concentration of 0.1g/L and one at 1.0g/L (our control, low and high copper concentrations, respectively). The plates were then incubated in a Biotek Synergy 2 spectrophotometer (Vermont, United States) for 24 hours at 37°C with an optical density (OD_600_) measurement taken every ten minutes.

### 16S rRNA amplicon sequencing

The 16S rRNA gene was amplicon sequenced in the three starting communities, as well as fifteen final communities from community A (5 of each copper treatment) and twelve final communities from both community B and C (4 of each copper treatment): not all replicates were sequenced due to budgetary constraints. DNA was extracted from the final communities using a DNeasy UltraClean Microbial Kit according to the standard protocol. The concentrations of DNA were then quantified using the Qubit HS DNA kit, before its purity was assessed using Nanodrop 260:280 ratios and its integrity by running samples on a gel (1% agarose). Sequencing was carried out at the University of Liverpool’s Centre for Genomic Research using an Illumina MiSeq. The raw fastq files were trimmed for the presence of Illumina sequences using ‘*Cutadapt*’ (v1.2.1; (39)). Sequence data was then processed and analysed in R using the ‘*dada2’* (40) and ‘*phyloseq’* (41) packages. After carrying out the standard full-stack workflow, we estimated error rates, inferred and merged sequences, constructed a sequence table, removed chimeric sequences and assigned taxonomy. The first 20bp of forward and reverse reads were trimmed during processing and all reads were truncated at 240bp. Taxonomies were assigned to amplicon sequence variants (ASVs) using the SILVA database (42). A phylogenetic tree was estimated using ‘*fasttree’* (43), allowing UniFrac distances between communities to be calculated (44). Sequences assigned to the phylum Cyanobacteria, as well as any reads that had not been assigned to at least the phylum level, were then removed. The community B inoculum was found to have 283 ASVs, however the inoculum of communities A and C both only contained two reads and so all three starting communities were removed from all further analysis. We then filtered the remaining samples to contain only ASVs with a total abundance >10, reducing the number of ASVs across passaged communities from 110 to 95. Samples had an average 16,220 reads, with a minimum of 8,950 and a maximum of 28,438 reads. The rarefaction curves of all samples plateaued, indicating the sequencing adequately represents the community, and sequencing depth was not associated with our treatments. As a consequence, we used the un-rarefied data for all downstream analyses.

### Statistical analysis

All analyses were carried out using R version 4.1.0 (45) and all figures were made using the *‘ggplot2’* package (46). For all analyses apart from the survival curves, we performed model selection to find the most parsimonious model by sequentially removing non-significant model terms (p>0.05) from linear models and nested models were compared using likelihood ratio tests. We used the *‘emmeans’* package (47) to compare differences between treatments when significant interactions occurred, and the false discovery rate (fdr) method to adjust p values for multiple testing (48). For each model, residual behaviour was checked using the *‘DHARMa’* package (49).

#### Analysing virulence of microbial communities

We tested for differences in virulence between treatments using Bayesian regression using the R package ‘*rstanarm*’ (50) to fit survival curves, and estimated parameters using the ‘*tidybayes*’ R package (51). A proportional hazards model with an M-splines baseline hazard was fitted, with copper, community, plus their interaction as fixed effects, and random intercepts for each sample to account for non-independency of observations (*i.e.* 20 *G. mellonella* were inoculated with the same sample). We used the Bayesian approach because it more easily handles random effects, allows us to easily calculate custom hazard ratios, and visualise uncertainty of the model compared with the more commonly used Cox method. Models used three chains with uninformative priors and were run for 3000 iterations. We assessed model convergence using Rhat values (all values were 1) and manually checked chain mixing. A second proportional hazards model was then run as before, but without the copper-community interaction term present. The leave-one-out (loo) validation technique, from the ‘*loo’* R package (52), was used to compare the two models.

For the model that best fitted our data (had the greatest loo weighting), log hazards were estimated for each copper treatment within each of the starting communities. In order to test for differences in virulence between copper treatments within communities, hazard ratios were calculated as the exponential of the difference between two log hazards. Median hazard ratios with 95% credible intervals (95%CI), and the probability that these hazard ratios were above 1 were then calculated. If the 95% credible intervals of the median hazard ratios did not cross 1, this was interpreted as a significant difference in virulence between two treatments. This model allowed us to test differences in virulence between copper treatments. In order to visualise variation in virulence across all communities, we additionally calculated the virulence of the individual replicates to test whether these traits were associated with virulence. To do this, we additionally calculated the hazard ratio of each final community compared to the average survival probability. We did this by estimating the log hazard for each of the 54 final communities using the random effect estimates, then comparing this value to the mean log hazard value of all of the 54 samples (*Mvirulence*), which was calculated by summing all of the hazard values and dividing the sum by the number of unique copper and community treatment combinations (*n*=54): 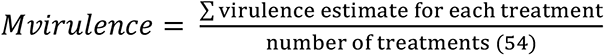 Consequently, samples with hazard ratios greater than one are more virulent compared to average, and samples with a value below one are less virulent compared to average.

#### Analysing the effect of copper on community productivity and copper susceptibility

The growth rates of whole passaged communities were first filtered to only include readings from 50 to 1080 minutes (0.83-18 hours), as the OD_600_ fluctuated randomly before and after these timepoints. Next, to control for any differences in starting optical density, either due to the inocula or to the different media, OD_600_ readings in any given well were corrected using OD_cor_ = OD_600_ – minimum OD_600_ in that well. The ‘*growthcurver*’ package (53) was then used to fit logistical growth curves and summarise the growth characteristics. For one community sample (community C, no copper treatment, when assayed with no copper), *growthcurver* was unable to converge and this sample was removed from downstream analysis (the community sample converged when assayed in both low and high copper media). *Growthcurver* provides estimates for three commonly-used growth traits: the exponential growth rate, *r*, the carrying capacity, *K*, and the area under the curve (AUC). AUC provides a metric that integrates information on *r*, *K*, and starting density, *N*_0_, and therefore is a metric of productivity (*i.e.* biomass accumulation over time). Therefore, we only compared the logistical area under the curves (AUC) across treatments. To test for differences in whole community AUCs, we used a linear model with AUC as the response variable, and copper, community and media, plus all interactions, as the explanatory variables. We then carried out pairwise comparisons to test the differences in AUC when grown in media containing no, low or high concentrations of copper: this allowed us to test whether passaged communities had different susceptibility to copper.

#### Community diversity and structure

We tested the effect of copper on the alpha diversity, evenness and composition of the final communities. Community evenness was calculated as Pielous’s evenness (Shannon index/log(number of observed ASVs)) (54). Alpha diversity, or richness, was taken as the total number of observed ASVs. We tested the effect of copper on evenness and alpha diversity using a linear model, with copper, starting community and their interaction as explanatory variables. Alpha diversity was log_10_ transformed to improve residual behaviour.

To calculate differences in the compositions of the communities, we used weighted UniFrac distances. UniFrac distance is preferable over other distance metrics because it incorporates phylogenetic distances between the ASVs observed within each community (44). As these distance values are dependent on the root of the phylogeny, we created a reproducible root by re-rooting to the longest branch that ended in a tip. To compare any differences in composition across copper treatments, we used the R packages ‘*phyloseq’* (41) and ‘*vegan’* (55). Specifically, we used ‘*vegan::adonis*’ to run a permutational ANOVA on UniFrac distance with 9999 permutations and ‘*vegan::betadisper*’ to analyse differences in group dispersion. Again, these models included copper, starting community and their interaction as explanatory variables. Permutational ANOVAs were simplified to the most parsimonious models as described above. UniFrac distances were used to carry out a principal coordinate analysis (PCoA).

In order to identify the ASVs that were affected by high copper concentrations (showed differences in abundance), we used the R package ‘*DESeq2*’ (56) to fit a negative binomial generalised linear model to the data. This method tests for significant differences in ASV abundances using Wald tests, and then uses the false discovery rate (fdr) method to correct for multiple testing. As the virulence of the low copper treatment did not significantly differ to the control and *DESeq2* only works with factors with two levels, only differences in ASV abundances between the high copper and control treatments were compared (n=83).

#### Associations between community virulence, productivity, diversity, and composition

Finally, to visually explore if there was an association between community-level virulence and community productivity, diversity or composition (PCoA1), we ranked each trait for all 54 samples. We ranked them because fitting statistical models to this data is problematic due to: 1) our values (particularly of PCoA1) not being evenly distributed across their range; 2) because it allows comparison of communities that have not been sequenced (15/54), and 3) because we have no prior expectations of how our predictors will affect virulence (e.g. in a linear or quadratic fashion etc.).

## Results

### High copper concentrations reduced the virulence of a wastewater community

Here, we tested the effect of copper on the virulence of three different sewage (influent) microbial communities (‘A’, ‘B’ and ‘C’) by incubating them with no copper, low (0.1g/L CuSO_4_) or high (1.0g/L CuSO_4_) copper concentrations. After passaging for three days with copper and one day without, we used the *G. mellonella* insect infection model to quantify the virulence of each community, with greater and/or faster mortality demonstrating higher virulence. The model containing the copper-community interaction better explained our data (it had a greater loo weighting than the model without the interaction: 0.574 to 0.426, respectively).This effect was due to differences in effect sizes, with the direction of the effect of the copper concentrations being consistent between the three starting communities (Fig. 1). Across all three influent communities, we found the high copper concentration to reduce virulence. In communities A and B this effect was large and significant, with 86.9% and 87.0% reductions in virulence compared to the no copper controls, respectively (community A: high vs control: 95%CI = 0.035-0.43; high vs low copper: 95%CI = 0.024-0.093; community B: high vs control: 95%CI = 0.035-0.486; high vs low copper: 95%CI = 0.047-0.734). In community C however, the 48.6% reduction in virulence compared to the control was not significant (high vs control: 95%CI = 0.136-1.95; high vs low copper: 95%CI = 0.071-1.10). In contrast, the low copper concentration had little effect on community virulence relative to the control (all hazard ratios between control and low copper communities encompassed 1: Community A: 95%CI = 0.426-4.61; Community B: 95%CI = 0.187-2.55; Community C: 95%CI = 0.486-6.72; Fig.1). These results demonstrate that high copper concentrations reduce virulence across bacterial communities, whereas low copper concentrations have little effect.

**Figure 1.**
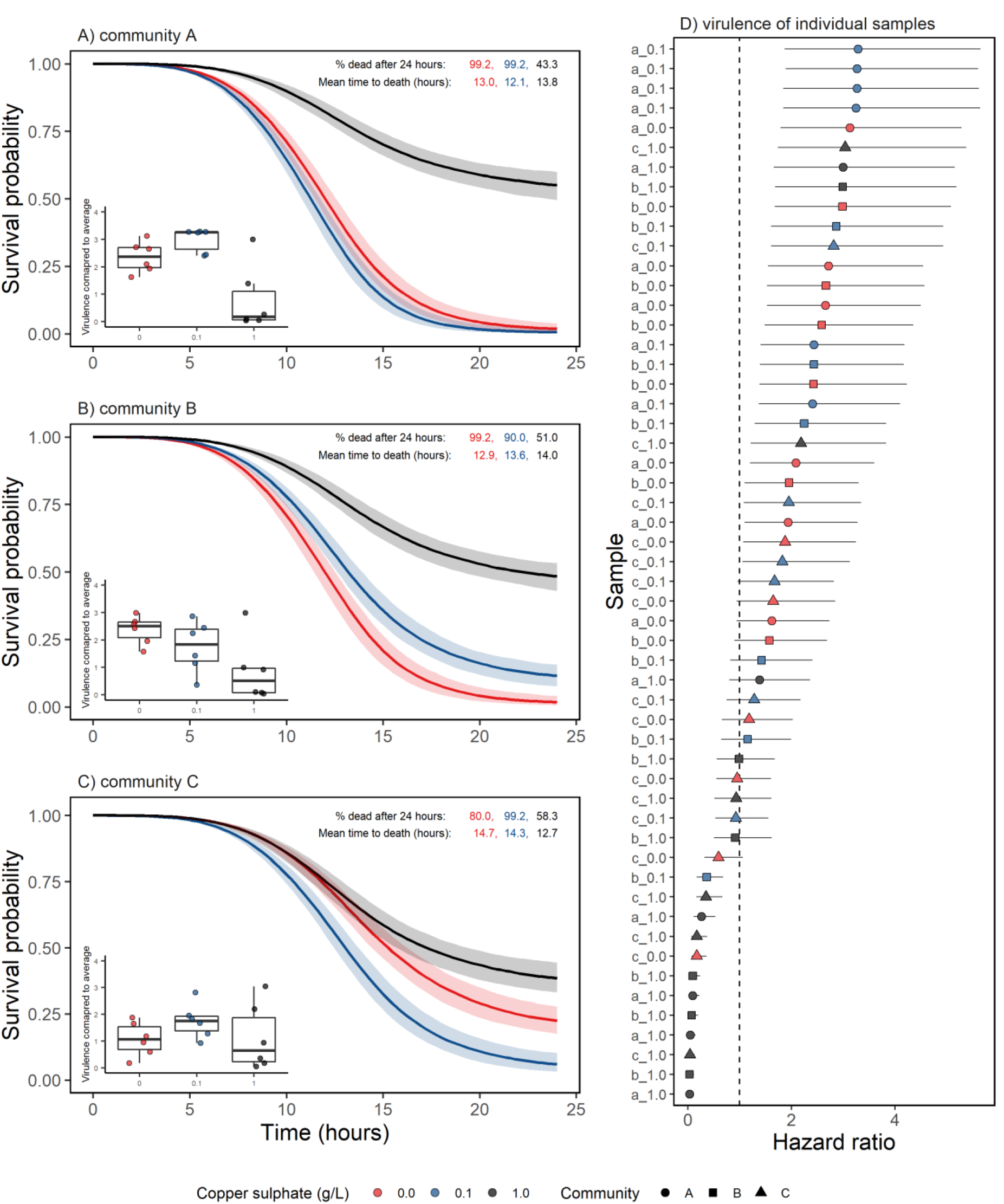
The survival probabilities for *Galleria mellonella* larvae injected with one of three communities (panels A-C) grown with no copper (red), low copper (0.1g/L CuSO_4_: blue) or high copper (1.0g/L CuSO_4_: black) for three days. Communities were grown for one day without copper before 20 larvae were injected per replicate (six replicates per treatment). Lines in panels A-C show median prediction and the shaded areas the 95% confidence interval. Inserted boxplots show the virulence (hazard ratio) of each final community compared to the average. Text in the top of panels A-C show the percentage of larvae that had died after 24 hours and the mean time in hours it took a larvae to die; numbers from left to right show the no copper control, low copper concentration and high copper concentration. Panel D shows a forest plot of each communities’ hazard ratio compared to average, with the horizontal lines showing the 95% confidence intervals. Samples to the left of the vertical line at x=1 in panel D are less virulent than average, whereas those to the right are more virulent than average. Sample labels on the y axis of panel D show the community followed by the copper concentration passaged in; community A is represented by circles, community B by squares and community C by triangles.

### Passaging with high copper concentrations reduced community productivity, and did not reduce susceptibility to high copper concentrations

To test whether exposure to copper affected the productivity of the communities and/or conferred increased copper resistance, we compared the productivity (AUC) of communities grown in the three different concentrations of copper (no, low, or high; Fig.2). Productivity was significantly affected by an interaction between copper history, and current copper growth conditions (copper history-media interaction: F_4,150_=5.10, p<0.001; Fig. 2). We found that when assayed in media containing no copper, productivity was significantly higher in communities that had been passaged with low copper concentrations (mean AUC = 589.7) than those passaged with no copper (mean AUC = 449.1; p=0.009) or high copper concentrations (mean AUC = 334.8; p<0.001), and that communities passaged with high copper concentrations were less productive than those passaged with low copper concentrations (p<0.019; Fig.2A). Consequently, we found that in the absence of copper, communities passaged with high copper had significantly lower productivity than those passaged with either no copper or low copper concentrations. Although community identity had a significant effect on productivity (community main effect: F_2,150_=9.65, p<0.001; Fig.2), the effects of copper selection regime (history) and current growth conditions were similar across different communities (copper history-community interaction: F_4,146_=1.47, p=0.214; media-community interaction: F_4,142_=0.69, p=0.600).

**Figure 2.**
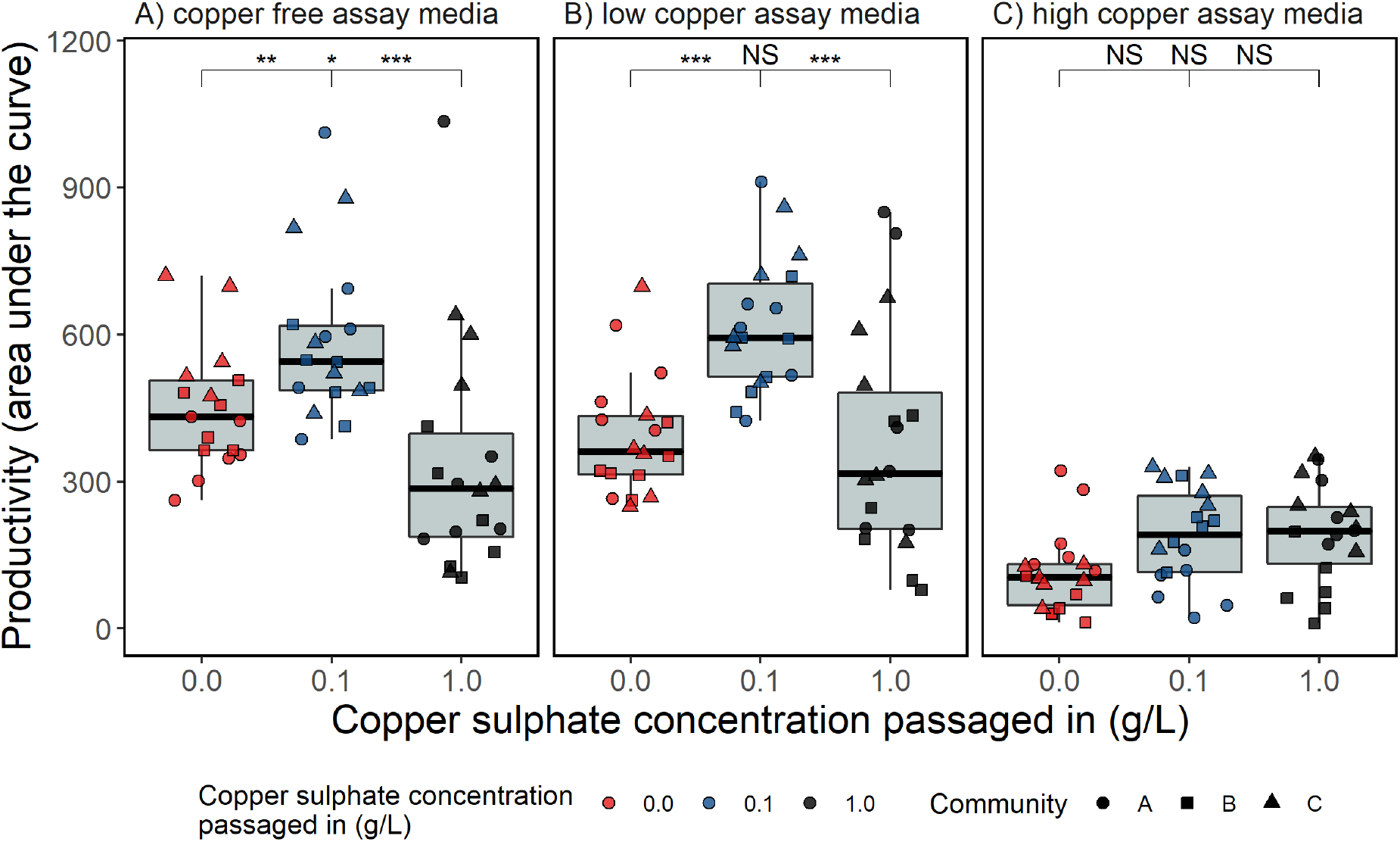
Area under the growth curves (AUCs) of three communities that have evolved with no copper (red shapes), low copper (blue shapes; 0.1g/L CuSO_4_), or high copper (black shapes; 1.0g/L CuSO_4_) for three days, and then for one day without copper. For this assay, each evolved community was grown for 18 hours in either assay media containing (A) no copper, (B) low copper (0.1g/L CuSO_4_) or (C) high copper (1.0g/L CuSO_4_) concentrations. The three communities are presented by different shapes: community A = circles, community B = squares and community C = triangles. Asterisks indicate significant differences between groups (*** = 0.001, ** = 0.01, * = 0.05, NS = non-significant), with the left value comparing the control to the low copper concentration, the middle value comparing the control to the high copper treatment and the right value comparing the low and high copper concentrations.

To explore if the reduction in productivity was due to a cost of copper resistance, we tested productivity in assay media containing our low and high copper concentrations. We found that when assayed in media containing low copper, the communities passaged with low copper remained the most productive, with the mean AUC value of 618.7 being significantly greater than the mean AUC value of 392.2 in the no copper treatment (p<0.001) and 379.3 in the high copper treatment (p<0.001). However, we now found productivity in the no copper and high copper concentration treatments to not be significantly different (p=0.790; Fig.2B). When assayed in media containing our high copper concentration, we found no significant difference in the productivity of any of our copper treatments (p=>0.177 for all contrasts; Fig.2C). However, we do note that productivity was very low when assayed in this media (mean AUC across all treatments = 165.5). These results therefore show that passaging in low copper concentrations is beneficial for productivity, but passaging in high copper concentrations is detrimental. Furthermore, they show that the reduced productivity of the communities passaged with high copper concentrations is not due to a cost of increased copper resistance, as they were no less susceptible to copper than the control or low copper treatments.

### Copper significantly reduces diversity and alters community composition

We tested the effect of copper on the evenness, alpha diversity and composition of a subset of passaged communities. The evenness (Pielou’s evenness) of the communities was not affected by either copper regime or community identity (copper main effect: F_2,33_=0.55, p=0.58; community main effect: F_2,33_=0.071, p=0.93; copper-community interaction: F_4,29_=1.24, p=0.32). However, alpha diversity (number of unique ASVs) was significantly affected by copper (copper main effect: F_2,34_=15.4, p<0.001; Fig. 3), but was not significantly different between communities (community main effect: F_2,34_=1.08, p=0.35; copper-community interaction F_4,30_=0.89, p=0.49). Alpha diversity in the high copper treatment (mean unique ASVs=14) was significantly lower than in either the control (high vs control: p<0.001) or the low copper treatment (low vs high p=0.009), which had on average 38 and 19 unique ASVs, respectively. However, low copper concentrations did not result in a significant difference in alpha diversity compared to the control (low vs control: p=0.061; Fig.3A). We note that the high copper treatment displayed a large variation in the number of unique ASVs compared to the low copper and control treatments, with a minimum of 1, a maximum of 50 and a median of 24 ASVs.

**Figure 3.**
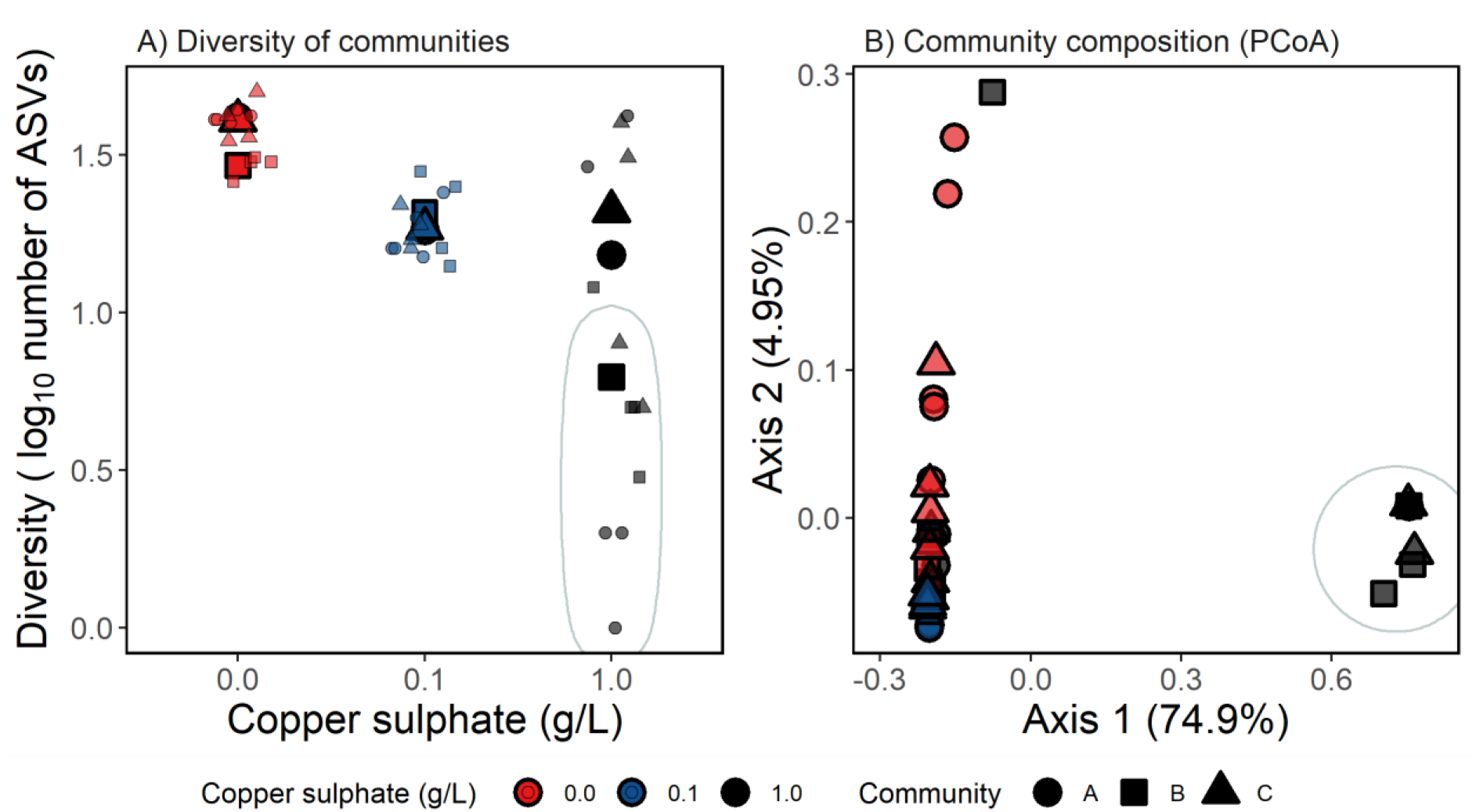
(A) Alpha diversity (log_10_ number of unique amplicon sequence variants) and (B) community composition (PCoA plot based on UniFrac distance) of three different complex microbial communites grown without copper (red), with low copper (0.1g/L CuSO_4_: blue) or with high copper (1.0g/L CuSO_4_: black) for three days, and then for one day without copper. Circles show community A, squares community B and triangles community C. In panel A, small shapes show individual replicates and larger shapes treatment means. The eight points circled in panel A represent the same communities as the points circled in panel B, demonstrating that the communites with fewest ASVs were of similar compositions; these were also the least productive communities (Fig.2A).

Similarly to alpha diversity, community composition (UniFrac distance) significantly differed between the copper treatments (copper main effect: F_2,34_=16.3, R^2^=0.48, p<0.001 Fig. 3B), but did not significantly differ between communities (community main effect: F_2,34_=0.77, R^2^=0.023, p=0.54; copper-community interaction F_4,30_=0.68, R^2^=0.042, p=0.64). However, in contrast to the alpha diversity, all copper treatments differed significantly in terms of their composition (p_adj_=<0.001 for all pairwise PERMANOVAs; Fig. 3B). The first principal coordinate (PCoA1) explained 74.9% of the variation in the composition, with eight of the thirteen (61.5%) communities from the high copper treatment having large PCoA1 values that made them very distinct from the other communities (circled points in Fig. 3B). These eight communities had the fewest ASVs and lowest productivity values, potentially explaining the bimodal distribution of AUC values seen in the high copper treatment (Fig.2D). The second principal coordinate explained only 4.95% of the remaining variation in composition, and generally separated the control and low copper treatments (Fig.3B).

### High copper concentrations significantly selected for *Leuconostoc lactis*, but against twenty other ASVs

To further explore changes in community structure, we examined which ASVs were selected for and against by the high copper concentration. To do this, we pooled the communities and focused on only the high copper treatment and the no copper control. Of the 83 unique ASVs found in either the high copper treatment or control, 21 significantly differed in abundance between the two treatments (a log2 fold change in abundance at a p value of <0.05). Of these 21 ASVs, we found only one ASV, identified as *Leuconostoc lactis*, to be positively selected for in the high copper treatment (log2 change in abundance= 7.91, p<0.001; Fig. 4). Although this taxon was found in all three starting communities sampled at the final time point, it only dominated high copper communities, was present in all eight of the circled points in figure 3, and made up >97.6% of five of the sequenced communities from the high copper treatment. These five communities had an average of only 9% *G. mellonella* mortality after 24 hours (mean mortality after 24 hours of all samples= 79.9%), with the one community consisting of 100% *L. lactis* having zero mortality after 24 hours. Although we found the abundance of 20 ASVs to be significantly reduced by the high copper concentration, three (two in the genus *Megasphaera*, and one identified as *Veillonella ratti*) were particularly affected, with log2 fold changes of -13.9, -15.0 and -16.0, respectively (Fig.4). These three ASVs however, never made up more than 0.25% of any community.

**Figure 4.**
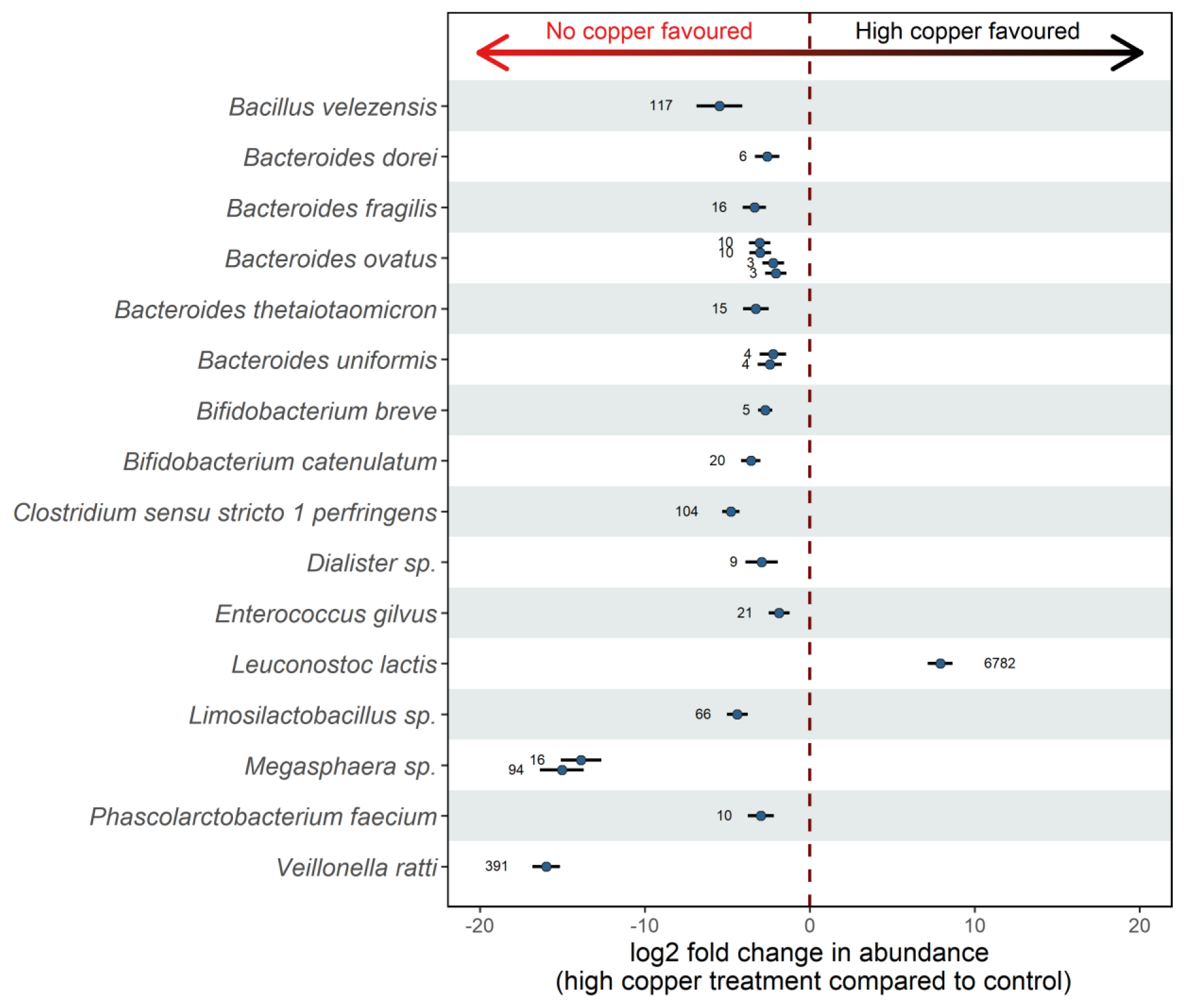
Amplicon sequence variants (ASVs) that differed significantly in abundance in the high copper versus the no copper treatment. ASVs are identified either to the genus or species level. Circles to the right of the vertical dashed line were positively selected for when grown with high copper, whereas those to the left were selected against. Numbers show the estimated mean number of that ASV present in the high copper samples. Only ASVs that were significantly affected after correcting for multiple testing using the fdr method (21 out of 83 ASVs; p<0.05) are shown. Solid lines show the 95% confidence intervals.

### Community virulence was associated with community composition

Finally, to visually explore which community trait was most associated with virulence, we ranked each community’s virulence (hazard value), productivity (AUC), diversity (the number of unique ASVs), and its composition (the first principal coordinate as this explained a large 74.9% of the variation in composition; PCoA1) (Fig.5). We found 10 of the 11 least virulent communities to be from the high copper treatment, and to have very low values of each of the traits (shown by purple boxes in Fig.5).

**Figure 5.**
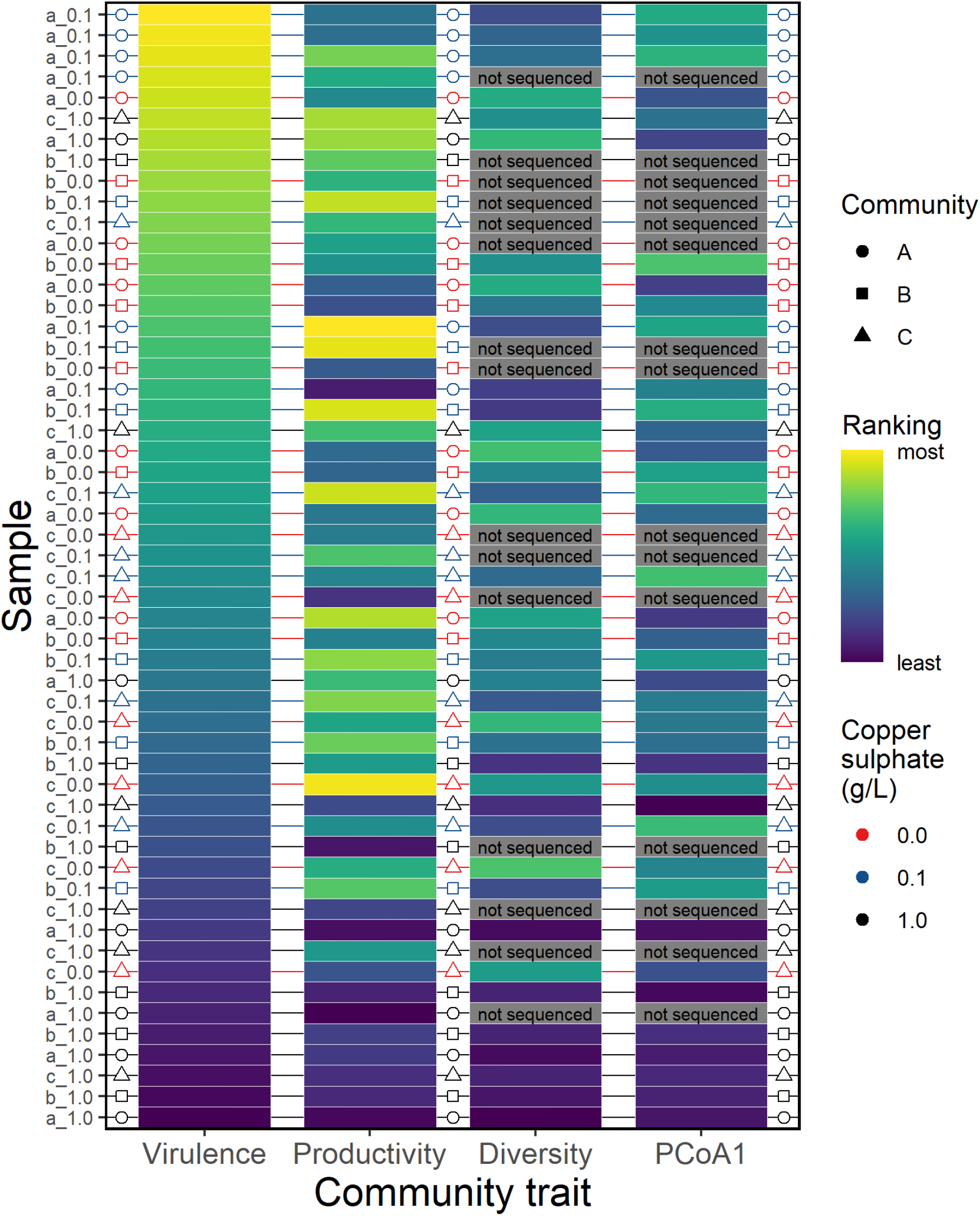
Ranked traits of 54 different microbial communites incubated either without copper, with low copper (0.1g/L CuSO_4_) or with high copper (1.0g/L CuSO_4_) for three days, and then for one day without copper. Communities are ordered by their virulence compared to one another in the first column. Column two shows their productivity (AUC) rank, column three their diversity (alpha diversity: number of unique ASVs) and column four a measure of community composition (-PCoA1). Yellow boxes show the communities with the highest value of that trait, and purple boxes the lowest value. Grey boxes show communities that were not sequenced (15 out of 54). Sample labels on the y axis show the community followed by the copper concentration passaged in.

## Discussion

Here, we tested the effect of environmentally relevant concentrations of copper on the virulence of three microbial communities taken from wastewater influent. We found that high copper concentrations reduced the virulence of all three communities, and for this reduction to be significant in two of the communities. However, low copper concentrations had little effect on virulence in any community. It has previously been shown that high concentrations of copper result in the altered composition of bacterial communities (57, 58), demonstrating the strong selective potential of metal pollution. Using 16S amplicon sequencing, we were able to show that the high copper treatment caused changes in community composition, and resulted in eight of the thirteen samples having severely reduced diversity with communities dominated by the same one or two genera. These same samples also had particularly low virulence. Moreover, we found that each of the three different starting communities had at least two final samples in this cluster of eight, demonstrating that this shift in composition was general across communities.

The association between high levels of copper and reduced virulence could potentially be explained by three distinct, but non-mutually exclusive, mechanisms. Firstly, a reduction in alpha diversity (species richness); secondly, reduced productivity (a low rate of biomass accumulation); and thirdly, selection for species with low virulence (identity effect). Below we discuss how each of these properties may be responsible for the observed decrease in virulence.

A reduction in species richness may decrease virulence by reducing the chance of species undertaking competition-mediated accelerated resource use within the host, as is frequently found in multi-species (‘polymicrobial’) infections (59–62). Moreover, it may also alter the expression of certain genes, including those associated with siderophore production, which can happen both in the presence of specific competing species and as community complexity increases (63). However, diversity is most plausibly likely to influence virulence by affecting the productivity of the community, as these traits are commonly found to be linked (27).

The reduction in community productivity associated with the high copper treatment may have reduced virulence for multiple reasons. First, and most likely, as greater growth requires more resources, more productive communities are likely to more efficiently exploit a host (29). Second, virulence factors produced by the community, such as toxins (64), may be produced in greater quantities by larger populations. In this respect, productivity of a community may offer an equivalent predictor of virulence as growth rate in single species infections (28–30).

Lastly, copper-imposed change in community composition may have a more direct consequence for virulence if the resulting community has a higher proportion of non-virulent species. For example, the eight sequenced high copper communities with the lowest virulence had almost identical compositions, each comprising fewer than ten unique ASVs, belonging to either the genus *Leuconostoc* or *Weissella*. Half of these communities were comprised solely of *L. lactis* – the only ASV to be selectively favoured under the high copper concentration. Whilst there is some evidence that this species can be pathogenic (65), it is widely regarded as non-pathogenic and is used in the dairy industry (66). Therefore, high copper may have reduced virulence by selecting against pathogenic species and favouring non-pathogenic species. Interestingly, previous work has found the supernatant from an isolate of *L. lactis* to have antimicrobial effects on several pathogenic species including *Staphylococcus aureus*, *Escherichia coli* O157, and *Proteusbacillus vulgaris* (67). Similarly, other members of the genera *Leuconostoc* and *Weissella* have been shown to have antimicrobial properties, most notably by reducing the formation of biofilms (68–70). It is therefore plausible that the high copper treatment exhibited lower virulence because it caused species sorting towards non-pathogenic species, and that this affect was possibly compounded by the resulting species potentially having antimicrobial traits.

As we find high copper to select for communities with reduced diversity, productivity, and with similar species identities, it is hard to tease apart which one has the largest effect on virulence. Furthermore, the virulence of whole communities – and the underlying traits – is not currently well understood, with little previous work to put our findings into context (but see (38) and (36)). Our recent work on bacterial pathogens associated with marine plastics is one of the few studies investigating community virulence (36). In that study, although community structure varied across communities with differential virulence, we found no relationship between virulence and diversity, nor did we explicitly test whether productivity, diversity or composition was the main driver explaining variation in virulence. Therefore, further work is required to experimentally disentangle their relative roles in community virulence. It is important to note when doing this, however, that these traits are likely inter-dependent. For example, previous work has shown a relationship between diversity and productivity, such that more diverse communities are generally more productive (27). Likewise, more diverse communities are more likely to contain virulent species, and so diversity may mask the identify effect.

Our finding that the low copper treatment did not affect virulence and that the high copper treatment reduced virulence is different from the previously observed positive association between copper and virulence in *Pseudomonas aeruginosa* (21). The difference between species-level and community-level results is most likely due to the main mechanism associated with both virulence and metal detoxification in *P. aeruginosa*, siderophore production, being likely to evolve differently in multi-species communities (24). Furthermore, it is expected that due to the short timescale of this current study species sorting is behind any observed changes in virulence rather than the evolution of metal detoxification mechanisms (71). We therefore highlight the importance of testing for the effects of a stressor at the community level, as they might differ from results based on single species experiments. Furthermore, as we find a non-significant effect of high copper on virulence in one of the three starting communities, we demonstrate that the response to an environmental stressor can differ across different communities.

In conclusion, our findings show that high concentrations of copper reduce the virulence of microbial communities. We suggest this can be explained by species sorting towards less diverse and less productive communities that contain a lower proportion of virulent species. As this study uses copper concentrations commonly found in copper polluted environments (31, 32), and microbial communities taken from wastewater, we reason that our findings may be applicable to environments impacted by human activity.

## Declarations

### Availability of data and materials

All data and code will be made publicly available upon acceptance for publication.

### Competing interests

The authors declare that they have no competing interests

### Funding

LL would like to thank the NERC FRESH GW4 award no. NE/R011524/1, DP the NERC IRF award NE/W008890/1, EH the UKRI Future Leaders Fellowship award MR/V022482/1, AB the NERC award NE/V012347/1, and SK and MV the NERC award NE/T008083/1.

## Notes

### Competing Interest Statement

The authors have declared no competing interest.

